# Genetic screening identifies glial adenosine metabolism and adenylate kinase 1 inhibition as a therapeutic target in alpha-synucleinopathy

**DOI:** 10.1101/2024.05.15.594309

**Authors:** MJ Sodders, E Okorie, H Tahmasebidehkordi, A Marathi, N Kumari, M Lakhani, A Schiro, M Shen, N Sriram, S Sarkar, AL Olsen

## Abstract

Neuronal α-synucleinopathies, including Parkinson’s disease (PD) and dementia with Lewy bodies (DLB), lack disease-modifying therapies. We developed a *Drosophila* model for identifying novel glial-based therapeutic targets for α-synucleinopathies. In this model, human α-synuclein is expressed in neurons and individual genes are independently knocked down in glia. We performed a forward genetic screen, knocking down the entire *Drosophila* kinome in glia in α-synuclein expressing flies. Among the top hits were five genes (*Ak1*, *Ak6*, *Adk1*, *Adk2*, and *awd*) involved in adenosine metabolism. Knockdown of each gene increased brain adenosine levels, improved locomotor dysfunction, reduced α-synuclein oligomer levels, and rescued neurodegeneration. We determined that the mechanism of neuroprotection requires adenosine signaling through the neuronal adenosine receptor (AdoR) in *Drosophila*. These experiments support glial adenosine as a novel therapeutic target in PD.

## Introduction

There are no disease-modifying therapies for Parkinson’s disease (PD) and dementia with Lewy bodies (DLB). Known PD and DLB risk genes are expressed in glia, and emerging evidence suggests that glia can contribute to non-cell-autonomous neurodegeneration[1–3]. The role of individual glial genes in pathogenesis is poorly understood, however. To identify novel glial-based therapeutic targets for α-synuclein, we developed a *Drosophila* α-synucleinopathy model [4] in which gene expression can be independently manipulated in neurons and glia[3,5] using the Q[6] and UAS-Gal4[7] expression systems (Figure 1A). In this model, wild type human α-synuclein is expressed in all neurons using the pan-neuronal driver *neuronal-synaptobrevin* (*nSyb-QF2*). These flies develop the key hallmarks of neuronal α-synucleinopathy, including death of cortical and dopaminergic neurons, motor impairment, and α-synuclein inclusions. In the glia, any gene of interest can be over-expressed or knocked down using the pan-glial driver *repo-Gal4*. To identify glial genes that either enhance or suppress neuronal α-synuclein toxicity, we performed a forward genetic screen, knocking down the entire *Drosophila* kinome (360 kinases) in glia in α-synuclein flies or control flies. Among the top hits identified in the kinome screen were five genes (*Ak1*, *Ak6*, *Adk1*, *Adk2*, and *awd*) involved in adenosine metabolism, a component of purine metabolism. Expression of purine metabolism genes is dysregulated in PD [8] and DLB[9] patient brains and in our *Drosophila* model[3,10]. The genes identified include two adenylate kinases (adenylate kinase 1 (*Ak1*) and adenylate kinase 6 (*Ak6*)), two adenosine kinases (adenosine kinase 1 (*Adk1*) and adenosine kinase 2 (*Adk2*)), and one nucleoside diphosphate kinase (abnormal wing discs (*awd)*).

**Figure 1:**
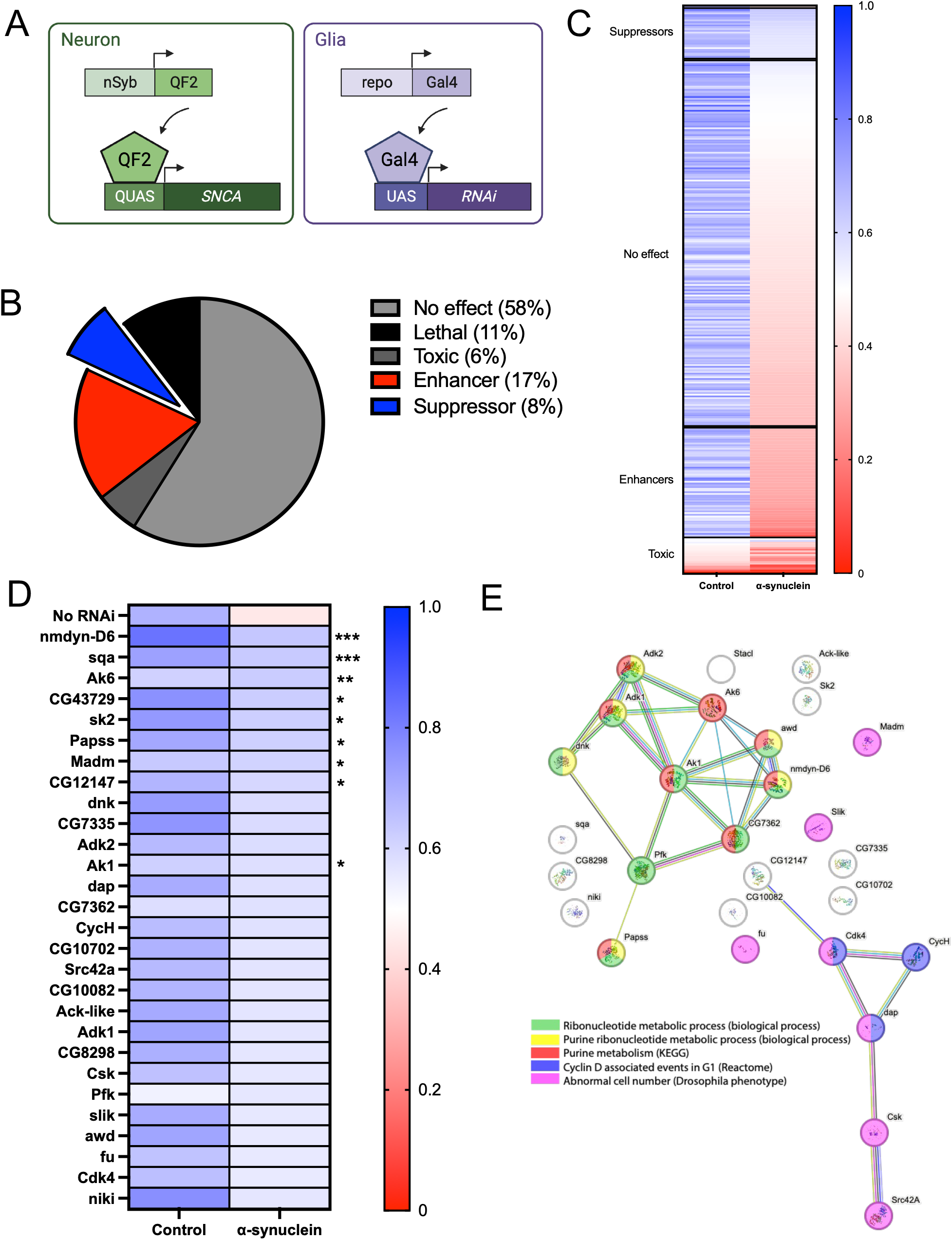
A kinome screen identifies purine metabolism as a glial therapeutic target pathway. A. Schematic of *Drosophila* model. Human α-synuclein is expressed in neurons, and genes are knocked down independently in glia. B. Percentage of lines characterized as suppressors, enhancers, toxic, lethal, or having no effect. C. Each line in the heatmap represents an individual kinase. D. Enlarged heatmaps demonstrating the suppressors. Statistical significance was determined by comparing the α-synuclein + RNAi of interest to the α-synuclein with no RNAi condition using two-way ANOVA with correction for multiple comparison testing with a false discovery rate of 0.5. *** = q-value <0.005, ** = q-value < 0.01, * = q-value < 0.05. q-values and original p-values for all lines are shown in Supplemental File 1. E. STRING analysis of the suppressors identifies two gene networks. Selected gene ontology and KEGG terms characterizing these genes are shown.

Glial knockdown of *Ak1*, *Ak6*, *Adk1*, *Adk2*, or *awd* resulted in increased adenosine and rescued neurodegeneration in *Drosophila*, suggesting that adenosine may be neuroprotective. Adenosine has many functions, at least two of which are germane to α-synucleinoapthies: 1) Secreted adenosine can signal through receptors, which have multiple, simultaneous, opposing effects. Adenosine signaling through the *inhibitory* A_1_ receptor *reduces* cAMP production and protects neurons from acute injury[11]. In contrast, adenosine signaling through the *excitatory* A_2A_ receptor *increases* cAMP production. The specific A_2A_ receptor antagonist istradefyelline is an FDA-approved symptomatic therapy for PD, supporting the relevance of this pathway in humans. 2) Adenosine can be metabolized to urate. Low urate is a PD [12–18] and DLB[19] biomarker, and urate is neuroprotective in models of PD[20–24], but a recent randomized clinical trial designed to increase urate levels failed to slow progression in PD[25]. Thus, while adenosine signaling and metabolism to urate are highly relevant to α-synucleinopathy, therapeutic targeting of this complex pathway has not been optimized.

To investigate the mechanism of rescue, we measured adenosine and its downstream metabolites after glial *Ak1*, *Ak6*, *Adk1*, *Adk2*, or *awd* knockdown. These experiments suggested that it is likely adenosine itself, as opposed to a downstream metabolite, that is responsible for rescue of neurodegeneration. We then determined that the neuronal adenosine receptor (AdoR) is required for rescue of neurodegeneration. In summary, we have identified adenosine metabolism as a novel potential glial-based therapeutic target for neuronal α-synucleinopathy.

## Results

We performed a forward genetic screen, knocking down the entire *Drosophila* kinome (360 kinases) in glia. The primary readout for the screen was a published behavioral assay that is based on locomotion[5]. Without any RNAi present, the average percentage locomotion in this assay for control flies was 67%, and the average percentage locomotion for α-synuclein flies was 44%. Based on results of the locomotion assay, genes were categorized as developmentally lethal, toxic, enhancer, suppressor, or having no effect (Figure 1B). Genes were categorized as developmentally lethal if no flies of the intended genotype eclosed. Suppressors were defined as those genes whose knockdown resulted in a 25% increase in locomotion in α-synuclein flies (Figure 1B-1D). Enhancers were defined as those genes whose knockdown results in a 25% decrease in locomotion in α-synuclein flies (Figure 1B-1C, Supplemental Figure 1). Toxic genes were defined as those whose knockdown results in a 25% decrease in locomotion in control flies. Any gene not meeting the 25% threshold was categorized as having no effect. A heatmap of the results for all non-lethal genes is shown in Figure 1C and all results are included as Supplemental File 1.

We performed STRING analysis (https://string-db.org, version 12.0) and found two networks among the suppressor genes (Figure 1E). No gene ontology or KEGG pathway analysis terms were statistically significantly enriched when comparing the list of 28 suppressors to the full list of kinases as a reference gene set, but, descriptively, one network consists primarily of purine metabolism related genes and the other of genes related to cell cycle and cell number (Figure 1E). STRING analysis of the enhancers showed one large network and one smaller network (Supplemental Figure 2). As with the suppressors, no gene ontology or KEGG pathway analysis terms were statistically significantly enriched when comparing to the full list of kinases as a reference gene set. Descriptively, most genes in the enhancer networks were involved in cell signaling, and specifically Wnt signaling (Supplemental Figure 2).

We are most interested in the suppressors identified in the kinome screen, as these genes might be able to rescue neurodegeneration if inhibited pharmacologically. Thus, we decided to focus further on the purine metabolism pathway, which has known relevance to α-synucleinopathy. Interestingly, among the 10 suppressor genes involved in purine metabolism identified in the kinome screen, 6 are involved specifically in adenosine metabolism, and their knockdown would be predicted to result in increased adenosine levels based on their predicted enzymatic function (Figure 2A). The genes include two adenosine kinases (*Adk1, Adk2*), two adenylate kinases (*Ak1, Ak6*), and two nucleoside diphosphate-kinases (*awd, nmdyn-D6*). Each has a human ortholog with high degree of homology (Figure 2B).

**Figure 2:**
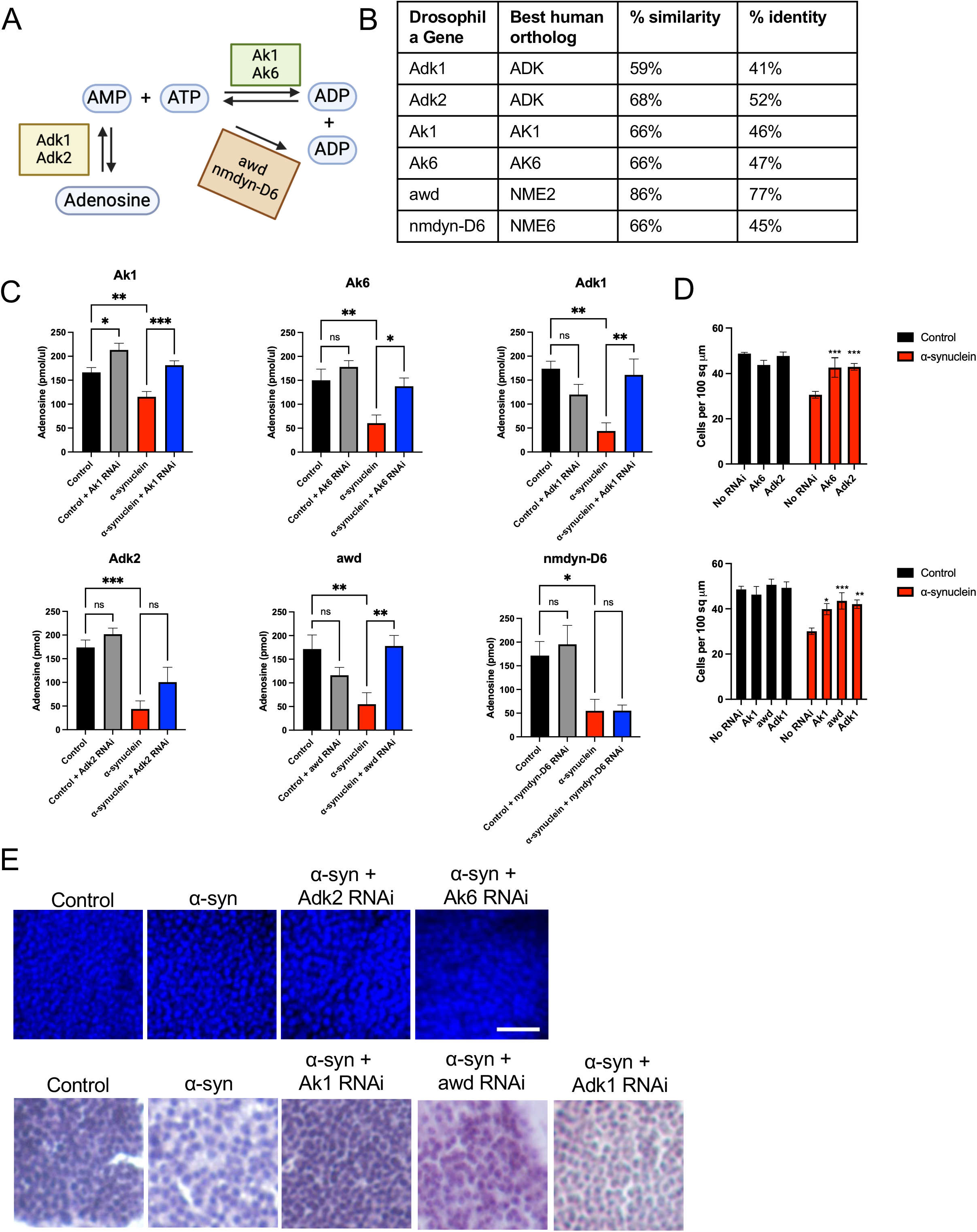
Glial knockdown of *Adk1, Adk2, Ak1, Ak6,* and *awd* increases adenosine and rescues neurodegeneration. A. Schematic of suppressor genes identified in the kinome screen whose knockdown is predicted to increase adenosine. B. Human orthologs of each gene. % identity and similarity to the human gene is indicated. C. Adenosine was measured using a fluorometric assay. Control flies are *nSyb-QF2, repo-Gal4/+* and α-synuclein flies are *nSyb-QF2, repo-Gal4, QUAS-*α*-synuclein/+*, each with or without a *UAS-RNAi*. n = 6 replicates of 2 heads per genotype. Statistical differences were determined by one-way ANOVA with Dunnett’s corrections for multiple comparison testing. ** = p <0.01, * = p <0.05. D. Cells were counted in the anterior medulla and normalized by area. n = 4-6 flies per genotype. Statistical significance was determined by comparing the α-synuclein + RNAi of interest to the α-synuclein with no RNAi condition using two-way ANOVA with Dunnett’s correction for multiple comparison testing. E. Representative images of the anterior medulla by whole mounted dissected brain stained with DAPI (top) or sectioned brain stained with hematoxylin (bottom). Scale = 10um. **** = p < 0.001, *** = p < 0.005, ** = p < 0.01, * = p < 0.05.

To determine whether knockdown of these 6 genes does increase adenosine, we used a fluorometric assay to measure adenosine levels in *Drosophila* heads (Figure 2C). We found statistically significantly increased adenosine after knockdown of *Ak1*, *Ak6, Adk1*, and *awd*. Knockdown of *Adk2* also led to increased adenosine, though the result was shy of statistical significance (p = 0.16). Knockdown of *nmdyn-D6* did not lead to increased adenosine. We therefore decided to focus the remainder of our analysis on the 5 genes (*Adk1, Adk2, Ak1, Ak6,* and *awd*) whose knockdown both increased adenosine and rescued locomotion in α-synuclein flies. We confirmed that knockdown of each gene rescued locomotion with a second independent RNAi line for 3 of the 5 genes in question, *Ak1, Ak2,* and *awd* (Supplemental Figure 3A). For *Adk1* knockdown, a second line caused toxicity in both control and α-synuclein flies (Supplemental Figure 3A), and for *Adk2*, a second line did not rescue locomotion (Supplemental Figure 3A), presumably due lack of efficient knockdown (Supplemental Figure 3B). Finally, we confirmed that knockdown of each gene rescues neurodegeneration by measuring total cell counts in the anterior medulla, an area of strong pathology in this model (Figure 2D). This was done by either immunofluorescence on whole mounted brains or immunohistochemistry on sectioned brains, based on the availability of equipment at the time of analysis (Figure 2E). Knockdown of nmdyn-D6, which did not increase adenosine as expected despite efficient knockdown (Supplemental Figure 3C), also did not rescue neurodegeneration (Supplemental Figure 3D). *nmdyn-D6* is expressed at low levels in the fly, so it may be that *awd* is the dominant nucleoside diphosphate kinase in the adenosine metabolism pathway (Supplemental Figure 3E). Collectively, these results provide strong support for adenosine as a novel therapeutic target pathway. Expressing a different pathogenic protein, the human amyloid beta 42 (Aβ42), in neurons had no effect on adenosine levels (Supplemental Figure 3F).

We next sought to determine whether the mechanism of rescue of neurodegeneration involves changes in α-synuclein. We have previously demonstrated that high molecular weight soluble α-synuclein oligomer levels are correlated with neurodegeneration in the *Drosophila* model[3]. Thus, we examined α-synuclein oligomer levels after knockdown of *Adk1, Adk2, Ak1, Ak2,* and *awd* in glia. We found that glia gene knockdown reduced both the absolute level of α-synuclein oligomers as well as the ratio of α-synuclein oligomers to α-synuclein monomer (Figure 3), demonstrating that increasing glial adenosine is having non-cell-autonomous effects on α-synuclein pathology.

**Figure 3:**
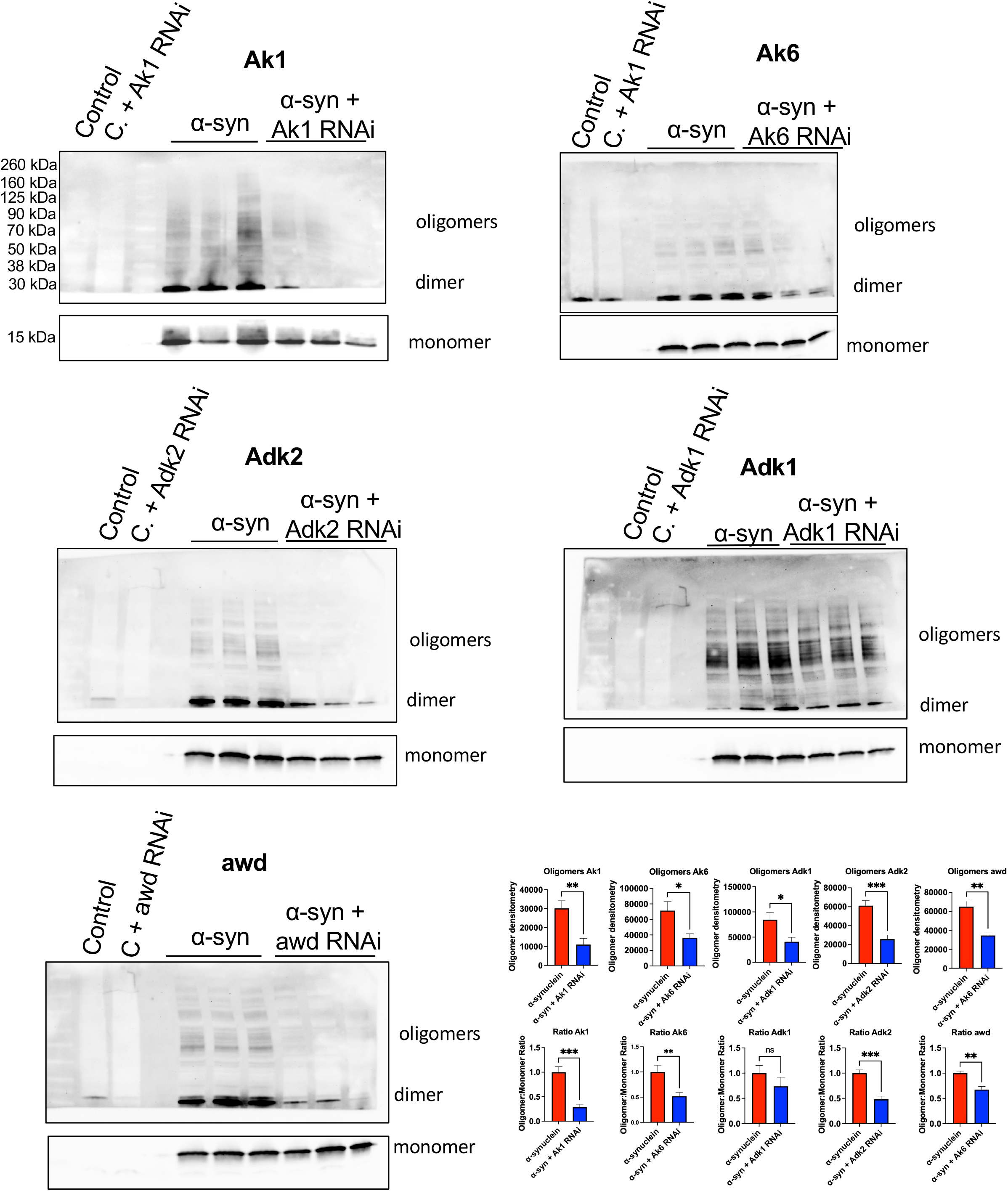
Glial knockdown of *Adk1, Adk2, Ak1, Ak6,* and *awd* reduces α-synuclein oligomer levels. Representative immunoblots stained with anti-α-synuclein clone 42 (1:5,000 mouse, BD Bioscience). Each immunoblot was cut to allow for longer exposure for the oligomers compared to the α-synuclein monomer. Oligomers were quantified from 2-3 independent experiments with 3 replicates per genotype. The total oligomer level and ratio of oligomer to monomer is shown on the bottom right. Statistical significance was determined by comparing the α-synuclein + RNAi of interest to the α-synuclein with no RNAi condition using Student’s t-test. *** = p < 0.005, ** = p < 0.01, * = p < 0.05.

Adenosine has many potential effects, including that it can enter the purine breakdown pathway. In humans and other closely related hominids[26], this pathway ends in metabolism to urate, whereas in nearly all other species, including *Drosophila*, urate is further metabolized to allantoin, and eventually to urea (Figure 4A). Many epidemiologic studies have found low serum or CSF urate levels in PD [12–18] and DLB[19], and urate has been shown to be neuroprotective in animal models of PD[20–24]. Thus, it is possible that the neuroprotective effects of adenosine are mediated not by adenosine itself, but rather by its metabolism to urate. To better understand the downstream metabolite changes that occur after increasing adenosine, we measured inosine, hypoxanthine/xanthine, and urate levels after knocking down each of the 5 genes (Figure 4B). These experiments identified more modest changes in downstream metabolites after glial knockdown of the 5 genes, all of which were smaller in amplitude than the changes in adenosine, as shown in the heatmap in Figure 4B.

**Figure 4:**
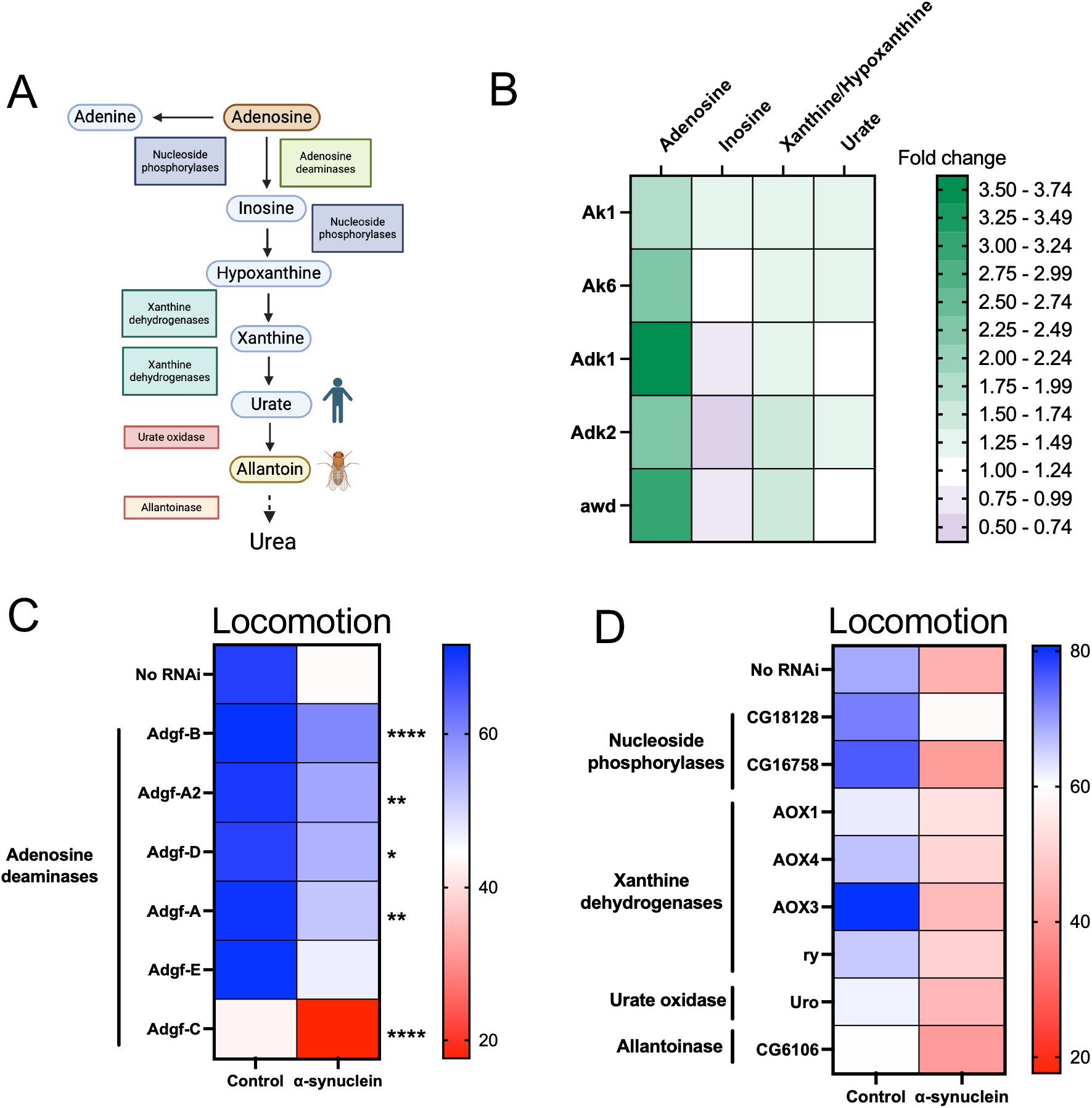
Rescue of neurodegeneration is likely mediated by adenosine rather downstream metabolites. A. Schematic of metabolites (ovals) and enzymes (boxes) involved in adenosine breakdown. Urate is the end-product of adenosine metabolism in humans. B. Heat map demonstrating relative change in adenosine metabolites after gene knockdown. C. Heatmap of locomotion after gene knockdown. Statistical significance was determined by comparing the α-synuclein + RNAi of interest to the α-synuclein with no RNAi condition using two-way ANOVA with correction for multiple comparison testing with a false discovery rate of 0.5. **** = q-value < 0.0001, *** = q-value <0.005, ** = q-value < 0.01, * = q-value < 0.05. D. Heatmap of locomotion after gene knockdown. No genes are statistically significant.

To further explore this, we identified all *Drosophila* genes involved in purine breakdown using the KEGG database (https://www.kegg.jp, release 109.1). We assessed locomotion in control and α-synuclein flies after individually knocking down all but 1 *Drosophila* gene involved in purine breakdown in glia (Figure 4A). One xanthine dehydrogenase, *AOX2*, was not able to be tested due to lack of any publicly available RNAi lines. First, we knocked down the 6 adenosine deaminases (*Adgf-A*, *Adgf-A2, Adgf-B, Adgf-C, Adgf-D, Adgf-E*), which convert adenosine to inosine (Figure 4A). With the notable exception of *Adgf-C* knockdown, which was toxic in both control and α-synuclein flies, knockdown of adenosine deaminases improved locomotion in α-synuclein flies (Figure 4C). In contrast, knockdown of genes further downstream in purine breakdown failed to improve locomotion in α-synuclein flies (Figure 4D). Thus, while we cannot exclude neuroprotective roles for inosine, hypoxanthine, xanthine, or urate, our data suggest that the effects are upstream of these metabolites.

Since the mechanism of neuroprotection likely involves adenosine itself, we then wondered whether adenosine receptor signaling is required. *Drosophila* have a single adenosine receptor, AdoR, which is expressed in both neurons and glia. We administered an adenosine receptor agonist, 2-chloroadenosine, in the *Drosophila* culture media, finding a dose-dependent improvement in locomotion in α-synuclein flies that peaked at a dose of 100 µM (Supplemental Figure 4). Based on these results, we selected a dose of 100 µM for further experiments. We then repeated the experiments with 100 µM 2-chloroadenosine, with knockdown of *AdoR* in either neurons or glia (Figure 5). Knocking down *AdoR* in glia did not affect the ability of 2-chloroadenosine to improve locomotion in α-synuclein flies (Figure 5A), whereas knocking down *AdoR* in neurons abolished the protective effect of 2-chloroadenosine (Figure 5B). These results demonstrate that adenosine receptor signaling in neurons is required for neuroprotection, identifying a non-cell-autonomous mechanism for glial adenosine.

**Figure 5:**
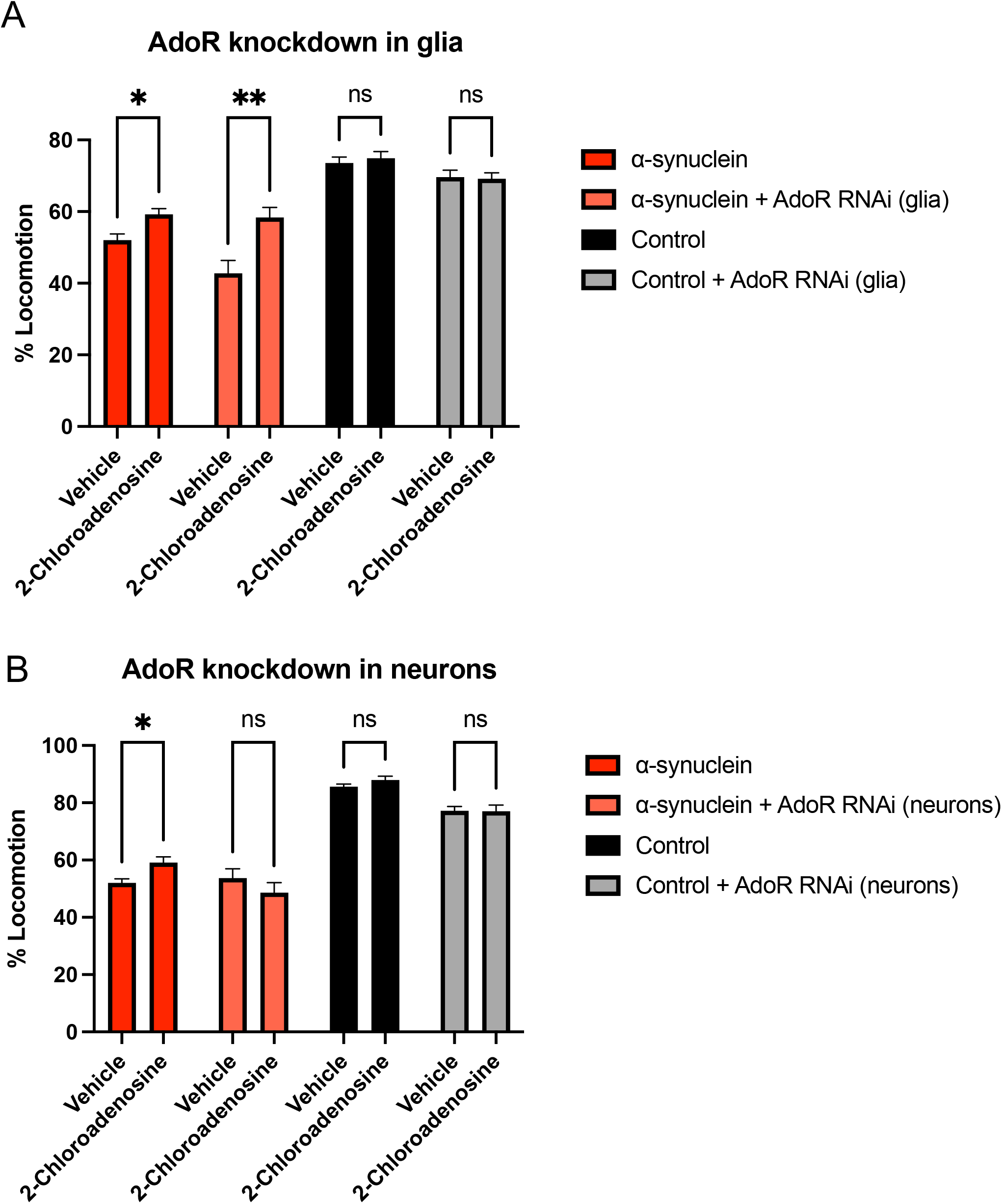
The neuronal adenosine receptor is required for neuroprotection. Flies were raised on culture media containing 100 µM 2-chloroadenosine or vehicle control (2% methanol) for 10 days post-eclosion. The locomotion assay was performed. n = minimum of 6 vials of at least 9 flies per genotype. Statistical significance was determined by comparing the α-synuclein + RNAi of interest to the α-synuclein with no RNAi condition using two-way ANOVA with Dunnett’s correction for multiple comparison testing. **** = p < 0.001, *** = p < 0.005, ** = p < 0.01, * = p < 0.05.

To further explore the downstream mechanisms of neuroprotection, we decided to focus on a single gene, adenylate kinase 1 (*Ak1*). We chose *Ak1* over the other genes because of attractive properties that make it an interesting potential therapeutic target. AK1 has been shown to be increased in both CSF and brain in human PD[8,27], suggesting that it may be pathologically elevated in the disease. Further, loss of *Ak1* appears to be relatively well-tolerated, as *Ak1*^-/-^ mice are viable, with only mild stress-induced phenotypes[28–33]. We have already shown that glial *Ak1* knockdown rescues neurodegeneration (Figure 2D). We next confirmed that this includes rescue of dopaminergic neuron loss (Figure 6A-B). Additionally, Ak1 knockdown reduces large α-synuclein aggregates (Figure 6C) and phosphorylated α-synuclein (Figure 6D), without affecting transgene expression (Figure 6D – see total α-synuclein monomer). Thus, glial *Ak1* knockdown rescues the two pathologic hallmarks of PD, dopaminergic neuron loss and Lewy-body like α-synuclein aggregates. The uncropped blot for total and phosphorylated α-synuclein is shown in Supplemental Figure 5.

**Figure 6:**
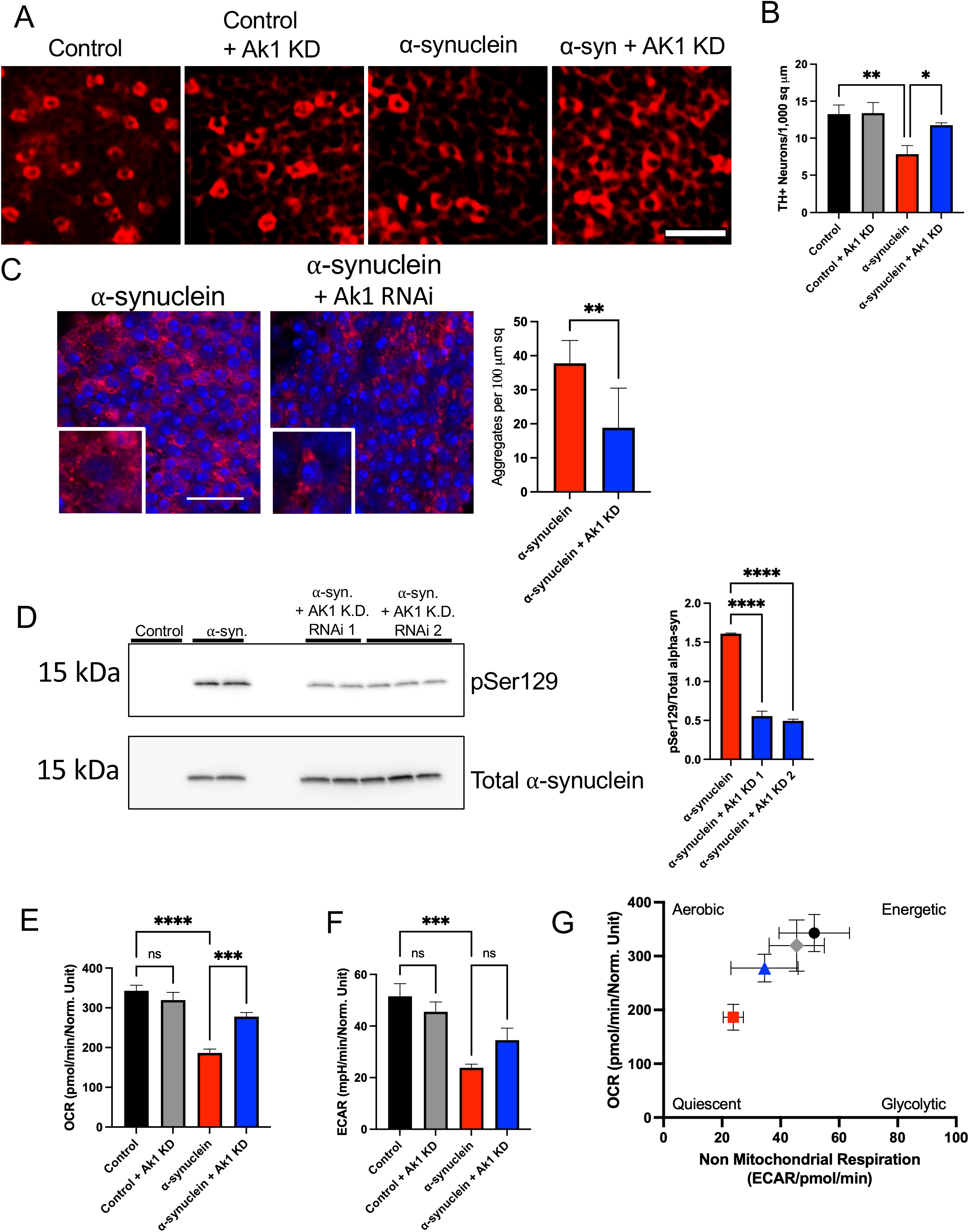
Glial Ak1 knockdown rescues dopaminergic neuron loss and pathologic α-synuclein. A. Dissected *Drosophila* brains were stained with tyrosine hydroxylase antibody (Immunostar, 1:200) and appropriate secondary. Scale bar = 10um. B. TH+ neurons in the anterior medulla were counted on 1 confocal slice at 20x magnification and normalized by area. n = 6 flies per genotype. C. Fixed tissue stained with anti-α-synuclein (clone 5G4, 1:50,000, mouse, Millipore) antibody that preferentially recognizes aggregated α-synuclein. Tissue is also stained with DAPI. Representative images are shown on the left. Scale bar = 10um. Aggregates were counted and normalized by area (right). n = 6 flies per genotype. D. Total and serine 129 phosphorylated α-synuclein was assessed by immunoblotting with α-synuclein H3C (1:10,000, mouse, Developmental Studies Hybridoma Bank), and phospho-serine 129 α-synuclein (1:5,000, rabbit, Abcam). Bands were quantified using Fiji (right). E. Oxygen consumption rate (OCR). Statistical differences were determined by two-way ANOVA with Dunnett’s corrections for multiple comparison testing. *** = p < 0.005, ** = p < 0.01, * = p < 0.05. F. Extracellular acidification rate (ECAR). Statistical differences were determined by two-way ANOVA with Dunnett’s corrections for multiple comparison testing. * = p < 0.05. G. The energetic profile is a way of visualizing the combined OCR and ECAR and indicates whether cells are more aerobic, glycolytic, quiescent or energetic. α-synuclein flies have a more quiescent profile than control flies due to decreased OCR and ECAR, and glial *Ak1* improves this. Error bars indicate SEM in the OCR and ECAR. n = 6 flies per genotype.

Ak1 is an adenylate kinase, part of a family of enzymes that catalyze the reversible reaction AMP + ATP 2 ADP. These enzymes are thought to sense the energy state of the cell and help coordinate a response by controlling adenine nucleotide levels[34]. We have previously demonstrated that α-synuclein flies have a quiescent bioenergetic profile, with impairments in both oxidative consumption rate (OCR) and extracellular acidification rate (ECAR), compared to control flies[10]. We show here that glial *Ak1* knockdown rescues the impairment in the OCR (Figure 6E), and partially rescues the impairment in the ECAR (Figure 6F), leading to an energetic profile similar to the control fly brain (Figure 6G).

In summary, through genetic screening, we identified glial adenosine metabolism as novel therapeutic target pathway in α-synuclein. We determined that knockdown of 5 genes (*Adk1, Adk2, Ak1, Ak6,* and *awd*) increases adenosine, reduces α-synuclein pathology, improves locomotion, and rescues neurodegeneration. Further, we identified glial *Ak1* as an especially promising target gene for rescuing loss of dopaminergic neurons and bioenergetic dysfunction.

## Discussion

We identified 5 genes (*Adk1, Adk2, Ak1, Ak6,* and *awd*) whose knockdown in glia results in increased adenosine, improvement in locomotion, reduction of α-synuclein pathology and rescue of neurodegeneration, suggesting that increasing adenosine is neuroprotective in α-synuclein flies. Of the 5 genes identified in the screen, we chose to focus further on *Ak1* both because of relevance to PD and because of properties that make it a potentially attractive therapeutic target. Both *Ak1* and *Ak6* are adenylate kinases. Their human orthologs are the identically named adenylate kinase 1 (*AK1)* and adenylate kinase 6 (*AK6)*, respectively. Relatively little is known about *AK6*, whereas increased *AK1* expression been associated with Alzheimer’s disease (AD) and PD in a handful of studies. Park et. al identified increased Ak1 expression in a mouse model of Alzheimer’s disease in whole brain lysates and further found that AK1 was increased in hippocampal neurons in human post-mortem AD[35]. AK1 expression was not examined specifically in glia, however[35]. Interestingly, another adenylate kinase family member, AK4, has also been associated with lower cognitive reserve[36] and faster cognitive decline[37] in AD. Further, Garcia-Esparcia et al reported that *AK1* mRNA was upregulated in frontal cortex in PD patients compared to controls[8]. In this study, the AK1 protein was detected in both neurons and astrocytes, but the cell type responsible for the increased mRNA expression was not investigated[8]. Finally, a recent proteomics study found that AK1 was increased in cerebrospinal fluid (CSF) in patients with PD compared to controls[27]. Despite its high degree of conservation and ubiquitous expression, systemic loss of Ak1 appears to be well-tolerated. *Ak1*^-/-^ mice are viable and fertile, with only mild exercise- or ischemia-induced phenotypes[28–33]. Glial *Ak1* knockdown may represent a “backdoor” to the adenosine metabolism pathway, that is, a way of altering adenosine levels indirectly, without attacking some of the most important enzymes involved in adenosine metabolism, thus avoiding some of the most critical side effects.

The next two genes we identified are *Adk1* and *Adk2*, which are adenosine kinases. They share a single human ortholog, adenosine kinase (*ADK*). ADK reduces adenosine levels by phosphorylating adenosine to AMP, and astrocytic ADK is the most important regulator of ambient adenosine levels[38]. Unfortunately, ADK is a poor therapeutic target: Genetic disruption of *Adk* in mice is lethal due to development of neonatal hepatic steatosis[39], missense mutations in humans cause hypermethioninemia, encephalopathy, and abnormal liver function[40], and initial attempts to inhibit ADK have led to on-target toxicity in the form of cerebral micro-hemorrhages[41]. Thus, preventing adenosine breakdown, at least by means of inhibiting ADK, is not likely to be a viable therapeutic strategy for increasing adenosine levels.

The final gene of interest, *awd*, is a nucleoside diphosphate kinase with many functions. Nucleoside diphosphate kinases transfer phosphates between nucleoside diphosphates and triphosphates. *awd* has high homology with both the human *NME1* gene and the *NME2* gene, as well as a readthrough product named *NME1-NME2*. *NME1*^-/-^ and *NME2*^-/-^ mice are viable, though *NME1*^-/-^ mice have growth retardation, and *NME1/NME2* double knockout mice die at birth from severe anemia[42]. NME1 and NME2 are suppressors of tumor metastasis[43]. *awd* also suppresses metastasis in *Drosophila*[44]. Garcia-Esparcia et al found reduced expression of NME1 in Parkinson’s disease substantia nigra[8]. Anatha et. al found that NME1 promoted neurite growth in SH-SY5Y cells and dopaminergic neurons[45], and *awd* is involved in synaptic vesicle recycling in *Drosophila*[46]. Collectively, these limited studies might argue in favor of increasing NME1 expression rather than reducing it and suggest further that *awd*/*NME1* could have different effects in neurons and glia.

One intriguing hypothesis to emerge from our results is that it may be adenosine itself rather than a downstream metabolite that is neuroprotective, as inhibiting downstream purine breakdown enzymes failed to rescue α-synuclein-induced impaired locomotion. Further exploring this topic is of critical importance in terms of clinical trial design, as attempts are underway to increase urate levels by giving patients dietary inosine in both PD and amyotrophic lateral sclerosis (ALS)[47]. Unfortunately, the phase 3 SURE-PD3 trial closed early due to an interim futility analysis, as clinical progression rates were not significantly different between participants who received inosine versus placebo[25]. Participants who received inosine in the trial had sustained increases in serum urate levels, and a previous phase 2 trial confirmed that dietary inosine also increases CSF urate levels[48], suggesting target engagement, although the differences were greater in women than men on subgroup analysis[49]. While there are many potential reasons for clinical trial failure, because adenosine is upstream of inosine, the inosine supplementation approach will not be successful if low urate levels in PD patients are merely a biomarker of adenosine dysfunction.

In summary, we have identified glial adenosine metabolism as a novel potential therapeutic target pathway in α-synucleinopathy. Further, we nominate Ak1 as a particularly attractive gene target for inhibition given that it is relevant to the human disease and loss of function appears well-tolerated.

## Materials and Methods

### Drosophila

All fly crosses and aging were performed at 25 °C. All experiments were performed at 10 days post-eclosion. All experiments include both male and female flies. All experiments include control (*nSyb-QF2, repo-Gal4/+*) and α-synuclein flies (*nSyb-QF2, repo-Gal4, QUAS-*α*-synuclein/+*), with or without a *UAS-RNAi*. The *nSyb-QF2 Drosophila* stock was provided by Dr. Christopher Potter. Transgenic RNAi stocks were obtained from the Bloomington *Drosophila* Stock Center or from Vienna *Drosophila* Resource Center. Two independent RNAi lines were used to confirm rescue of locomotor impairment following *Ak1*, *Ak6*, or *awd* knockdown: *UAS-Ak1 RNAi TRiP.HMC03355*, *UAS-Ak1 RNAi TRiP.GL00177*, *UAS-Ak6 RNAi TRiP.GL00285, UAS-Ak6 RNAi GD14036*, *UAS-awd RNAi TRiP.GL00013*, *UAS-awd RNAi TRiP.HMJ02099*. The first line listed in each pair was used for all other experiments involving gene knockdown, with the exception that *UAS-Ak6 RNAi GD14036* was used for the α-synuclein oligomer experiments (Figure 3). For the *Adk1* knockdown, only a single RNAi line (*UAS-Adk1 RNAi TRiP.HMC06361*) was used, because a second RNAi line failed to replicate the locomotion result (*UAS-Adk1 RNAi KK106704*), likely due to an off-target toxic effect, as the line was toxic in both control and α-synuclein flies (Supplemental Figure 3A). For the *Adk2* knockdown, only a single RNAi line (*UAS-Adk2 RNAi TRiP.GL00036*) was used, because the second line (*UAS-Adk2 RNAi TRiP.HMC05509*) did not significantly rescue locomotion (Supplemental Figure 3A), likely due to failure of knockdown (Supplemental Figure 3B). Efficiency of knockdown was validated using the systemic da-Gal4 driver. The QUAS-Abeta42 line used in Supplemental Figure 3F is P{QUAS-Abeta42.l}7 and was created as a model of Alzheimer’s disease[50]. It was obtained from the Bloomington *Drosophila* Stock Center (stock 83348).

### Kinome screen

A list of all *Drosophila* kinases was obtained from flybase.org. Every fly kinase tested in the screen has at least one human ortholog. One transgenic UAS-RNAi line for each kinase was purchased from the Bloomington *Drosophila* Stock Center. Control or α-synuclein flies were crossed to each transgenic UAS-RNAi line. If no progeny of the intended genotype eclosed in 3 independent crosses, the UAS-RNAi line in question was considered developmentally lethal (Figure 1B). For all other crosses, locomotion was measured in control or α-synuclein flies harboring the UAS-RNAi, using a previously published assay[5]. Results were compared to those of control or α-synuclein flies without any RNAi. Each line was categorized as having no effect, being a suppressor, being an enhancer, or being toxic, based on the degree of change in locomotion with the UAS-RNAi line compared to the control or α-synuclein flies without the RNAi. Suppressors were defined as RNAi lines that caused 25% improvement in locomotion in α-synuclein flies compared to the α-synuclein baseline. Enhancers were defined as RNAi lines that caused 25% decrement in locomotion in α-synuclein flies compared to the α-synuclein baseline. Toxic lines were defined as lines that caused 25% decrement in both α-synuclein flies and control flies compared to their respective baselines. All other results were classified as having no effect. Raw data for the kinome screen are included in Supplemental File 1.

### Adenosine, inosine, hypoxanthine/xanthine, and urate assays

Flies were anesthetized, then snap frozen on dry ice to halt metabolic reactions. After freezing, heads were removed and homogenized in assay buffer (Abcam). Homogenates were deproteinized using the Deproteinizing Sample Preparation Kit - TCA (Abcam ab204708). Fluorometric kits for adenosine (Abcam ab211094), inosine (Abcam ab126286), hypoxanthine/xanthine (Abcam ab155900), and urate (Abcam ab65344) were used per the manufacturer’s instructions. The hypoxanthine/xanthine kit does not distinguish between hypoxanthine and xanthine. Each experiment included 6 replicates per genotype, with 1 head per replicate for hypoxanthine/xanthine and 2 heads per replicate for the other assays.

### Quantification of total cell counts

For knockdown of *Ak1*, *Adk1*, and *awd,* total cells were counted as previously described[51]. Briefly, flies were fixed in formalin (Fisher Scientific SF9804) and embedded in paraffin (Fisher Scientific 83-30). 2 μm serial frontal sections were prepared through the entire fly brain. Slides were processed through xylene (Fisher Scientific X3P-1GAL), ethanols (Fisher Scientific 62-15), and into water, then stained with hematoxylin (Fisher Scientific 22-050-206). Images of the anterior medulla from hematoxylin-stained formalin-fixed paraffin-embedded tissue were captured at 40X magnification using brightfield microscopy. One image per fly and 6 flies per genotype were used for quantification. The nuclei in each tissue section were counted and normalized to the area of the section. dx.doi.org/10.17504/protocols.io.4r3l24on4g1y/v1 For knockdown of *Ak6* and *Adk2*, cells were counted using a modified whole-mount protocol[52]. Briefly, fly brains were dissected, fixed in 4% paraformaldehyde (Fisher AA47392-9M), permeabilized, stained with DAPI (Thermo Fisher D1306), and whole mounted with Slowfade Diamond mounting media (Fisher Scientific S36967). Images of the anterior medulla were captured by confocal microscopy at 20X magnification. One image per fly and 6 flies per genotype were used for quantification. The nuclei were counted and normalized to the area of the section using Fiji version 2.3.0/1.53q.

### Quantification of TH+ neuron counts

Fly brains were dissected, fixed in 4% paraformaldehyde, permeabilized, stained with DAPI, and stained with primary anti-tyrosine hydroxylase antibody overnight (1:200, mouse, Immunostar 22941). The next day brains were stained with appropriate secondary antibody and whole mounted. Images of the anterior medulla were captured by confocal microscopy at 20X magnification. One image per fly and 6 flies per genotype were used for quantification. Images were processed using Fiji version 2.3.0/1.53q. The number of TH+ neurons was normalized to the area of the section.

### Quantification of α-synuclein aggregates

α-synuclein aggregates were quantified as previously described[51]. Briefly, flies were fixed in formalin (Fisher Scientific SF9804) and embedded in paraffin (Fisher Scientific 83-30). 4 μm serial frontal sections were prepared through the entire fly brain. Slides were processed through xylene (Fisher Scientific X3P-1GAL), ethanols (Fisher Scientific 62-15), and into water. Microwave antigen retrieval with 10 mM sodium citrate (Fisher Scientific S279-3), pH 6.0, was performed. Slides were blocked in 2% milk (Millipore Sigma M7409) in PBS (Fisher Scientific BP665-1) with 0.3% triton X-100 (Thermo Scientific AAA16046AP) for 1 hour then incubated with anti-α-synuclein 5G4 antibody (1:50,000, mouse, Millipore MABN389) in 2% milk in PBS with 0.3% triton X-100 at room temperature overnight. Slides were incubated with fluorophore-conjugated secondary antibody in 2% milk in PBS with 0.3% triton X-100 for 1 hour (1:200, Alexa 555, Invitrogen A-21424) then mounted with DAPI-containing Fluoromount medium (Southern Biotech 0100-20). Immunofluorescence microscopy was performed on a Zeiss LSM 800 confocal microscope. Images of the anterior medulla from immunofluorescence-stained formalin-fixed paraffin-embedded tissue were captured on a confocal microscope at 63X magnification. One image per fly and 6 flies per genotype were used for quantification. The α-synuclein aggregates in each tissue section were counted and normalized to the area of the section. Images were processed using Fiji version 2.3.0/1.53q. dx.doi.org/10.17504/protocols.io.81wgby61nvpk/v2

### Western blotting of fly heads

Fly heads were homogenized in 2X Laemmli buffer (Sigma Aldrich S3401), boiled for 10 minutes, and centrifuged. SDS-PAGE was performed (Bio-Rad) followed by transfer to 0.2 µm nitrocellulose membrane (Bio-Rad 1704270) and microwave antigen retrieval in PBS. Membranes were blocked in 2% milk in PBS with 0.05% Tween-20 (Fisher PI85114) for 1 hour, then immunoblotted with appropriate primary antibody in 2% milk in PBS with 0.05% Tween-20 overnight at 4 °C. Primary antibodies used include α-synuclein H3C (1:10,000 to 1:100,000, mouse, Developmental Studies Hybridoma Bank H3C) and phospho-serine 129 α-synuclein (1:5,000, rabbit, Abcam ab51253). Membranes were incubated with appropriate horseradish peroxidase-conjugated secondary antibodies (1:50,000) in 2% milk in PBS with 0.05% Tween-20 for 3 hours. Signal was developed with enhanced chemiluminescence (Thermo Fisher 32106). dx.doi.org/10.17504/protocols.io.8epv5j96jl1b/v2

### Oligomer assay

As previously published[3], 20 fly heads per genotype were homogenized in 20 ul TNE lysis buffer (10 mm Tris HCl, 150 mM NaCl, 5 mM EGTA, 0.5% nonidet-40) supplemented with HALT protease and phosphatase inhibitor (Fisher Scientific 78440). The homogenate was briefly spun down to remove debris. The remaining supernatant was ultracentrifuged at 100,000 x g for 1 hour at 4° C. The supernatant was transferred to a new tube and combined with 2X Laemmli buffer (Sigma Aldrich S3401) at a 1:1 ratio. SDS-PAGE was then performed as above, except without boiling samples and without microwave antigen retrieval. Primary antibody used was α-synuclein clone 42 (1:5,000 mouse, BD Bioscience), and secondary antibody used was the appropriate horse radish peroxidase-conjugated antibody (1:20,000) for 1 hour.

### Locomotion assay

Adult flies were aged in vials containing 9-20 flies per vial. At day 10 post-eclosion, flies were transferred to a clean vial (without food) and given 1 minute to acclimate to the new vial. The vial was then gently tapped three times to trigger the startle-induced locomotion response, then placed on its side for 15 seconds. The percentage of flies still in motion was then recorded. Differences between genotypes were measured and statistical significance assessed by two-way ANOVA using GraphPad Prism version 10.2.3.dx.doi.org/10.17504/protocols.io.4r3l226p4l1y/v1

### Seahorse assay

The oxygen consumption rate (OCR) and extracellular acidification rate (ECAR) were measured using a Seahorse XFe96 metabolic analyzer. Brains were dissected and plated at one brain per well on Seahorse XFe96 flux pack plates (Agilent 103793-100) following the manufacturer’s protocol and as previously published[10]. The OCR values were normalized to DNA content using a CyQUANT assay (ThermoFisher C7026) following the manufacturer’s protocol.

### Quantitative real-time qPCR

Gene knockdown was confirmed using qRT-PCR. *Drosophila* primers were chosen from the DRSC FlyPrimerBank [53]. 6 heads per genotype were used. Total RNA was prepared with Qiazol (Qiagen 79306) per the manufacturer’s instructions then treated with DNase (Thermo Fisher EN0521) for 15 minutes. cDNA was prepared using a High Capacity cDNA Reverse Transcription Kit (Applied Biosystems 43-688-14), then amplified with SYBR green (Bio-Rad 1708882) on a StepOne Plus Real-Time PCR system (Applied Biosystems 4376600). Relative expression was determined using the ΔΔCt method, with normalization to *RPL32* used as a housekeeping gene for *Drosophila*, and normalization to GAPDH for human cells. dx.doi.org/10.17504/protocols.io.q26g7y4r1gwz/v1

### 2-chloroadenosine

2-Chloroadenosine was purchased from Sigma and reconstituted in methanol to a stock concentration of 100 mM. The stock was further diluted in water to the final concentration (1 µM to 1 mM) and mixed at a 1:1 ratio with *Drosophila instant* culture media (Carolina Formula 4-24). 2% methanol served as a vehicle control. Flies were reared on treated culture media from the time of eclosion to day 10 post-eclosion and flipped to new treated media every 2-3 days. dx.doi.org/10.17504/protocols.io.5jyl8j9n9g2w/v1

### Statistics

All statistical analysis was performed using GraphPad Prism version 10.1.

### Data availability

The data that support the findings of this study are available from the corresponding author upon request. The supplemental files associated with this manuscript have been deposited in Zenodo: https://doi.org/10.5281/zenodo.11370503. The data, code, protocols, and key lab materials used and generated in this study are listed in a Key Resource Table alongside their persistent identifiers at https://zenodo.org/records/13623853. An earlier version of this manuscript was posted to biorxiv on 5/15/24 at doi: https://doi.org/10.1101/2024.05.15.594309.

## Acknowledgments

The research was funded in part by Aligning Science Across Parkinson’s [ASAP-000301] through the Michael J. Fox Foundation for Parkinson’s Research (MJFF). For the purpose of open access, the author has applied a CC BY public copyright license to all All Author Accepted Manuscripts arising from this submission.

Other sources of funding include K08NS109344 (Olsen), 5P30AG062421 (Olsen), 2021A003323 the George C. Cotzias Award from the American Parkinson’s Disease Association (Olsen), R00ES033723 (Sarkar), P30ES001247 (Sarkar), and AWD00007930 Alzheimer’s Association Research Fellowship to Promote Diversity (Kumari).

The HC3 α-synuclein antibody was provided by the Developmental Studies Hybridoma Bank, created by the NICHD of the NIH and maintained at The University of Iowa, Department of Biology, Iowa City, IA 52242. Drosophila stocks obtained from the Bloomington Drosophila Stock Center (NIH P40OD018537) were used in this study. We thank the Transgenic RNAi Project (TRiP) at the Harvard Medical School (NIH-NIGMS R01GM084947) for making transgenic RNAi stocks.

## Supplemental Figure Legends

Supplemental Figure 1: Heatmap of enhancers from the kinome screen. Statistical significance was determined by comparing the α-synuclein + RNAi of interest to the α-synuclein with no RNAi condition using two-way ANOVA with correction for multiple comparison testing with a false discovery rate of 0.5. *** = q-value <0.005, ** = q-value < 0.01, * = q-value < 0.05. q-values and original p-values for all lines are shown in Supplemental File 1.

Supplemental Figure 2: STRING network analysis of enhancers from the kinome screen. Selected gene ontology and KEGG terms characterizing these genes are shown.

Supplemental Figure 3: A. Locomotion was measured with an independent RNAi-line. n = 6 vials per genotype of 9-20 flies per vial. Statistical significance was determined by comparing the α-synuclein + RNAi of interest to the α-synuclein with no RNAi condition using two-way ANOVA with Dunnett’s correction for multiple comparison testing. **** = p < 0.001, *** = p < 0.005, ** = p < 0.01, * = p < 0.05. B. Validation of gene knockdown for all RNAi lines. N = 6 heads per genotype. Expression of each gene in control was normalized to a level of 1, and statistical significance was determined by one sample t-test comparing the expression of each gene after knockdown to a theoretical mean of 1. Efficiency of knockdown was validated by crossing each RNAi line to *da-Gal4* driver. C. Validation of gene knockdown for the nmdyn-D6 line. D. nmdyn-D6 knockdown did not affect neurodegeneration. E. nmdyn-D6 expression is low in *Drosophila* compared to expression of *awd*. The image shown was obtained from flybase.org using publicly available data from the Fly Cell Atlas Project[54]. F. Expression of QUAS-Abeta42 in neurons does not affect adenosine. N = 6 heads per genotype.

Supplemental Figure 4: Flies were raised from the time of eclosion to day 10 post-eclosion on media supplemented with 2-chloroadenosine or vehicle (2% methanol) control. The locomotion assay was performed at day 10. N = minimum 6 vials of 11-20 flies per vial.

Supplemental Figure 5: The blots that are cropped in Figure 6 are shown here in full. Supplemental File 1: Complete results of kinome screen.

