## Supplemental Figures for "Genetic screening identifies glial adenosine metabolism and adenylate kinase 1 inhibition as a therapeutic target in alpha-synucleinopathy"

### Slide 1
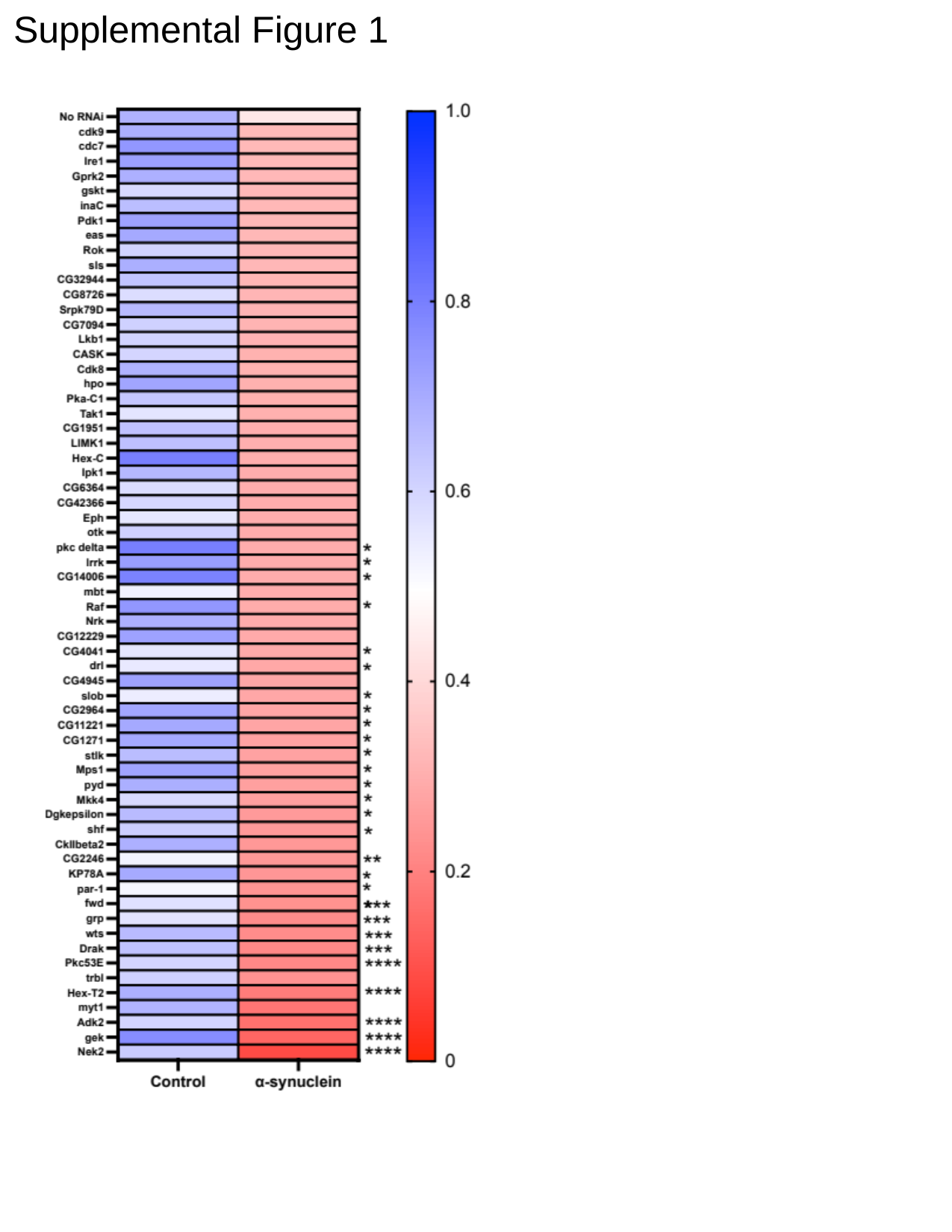

Supplemental Figure 1

### Slide 2
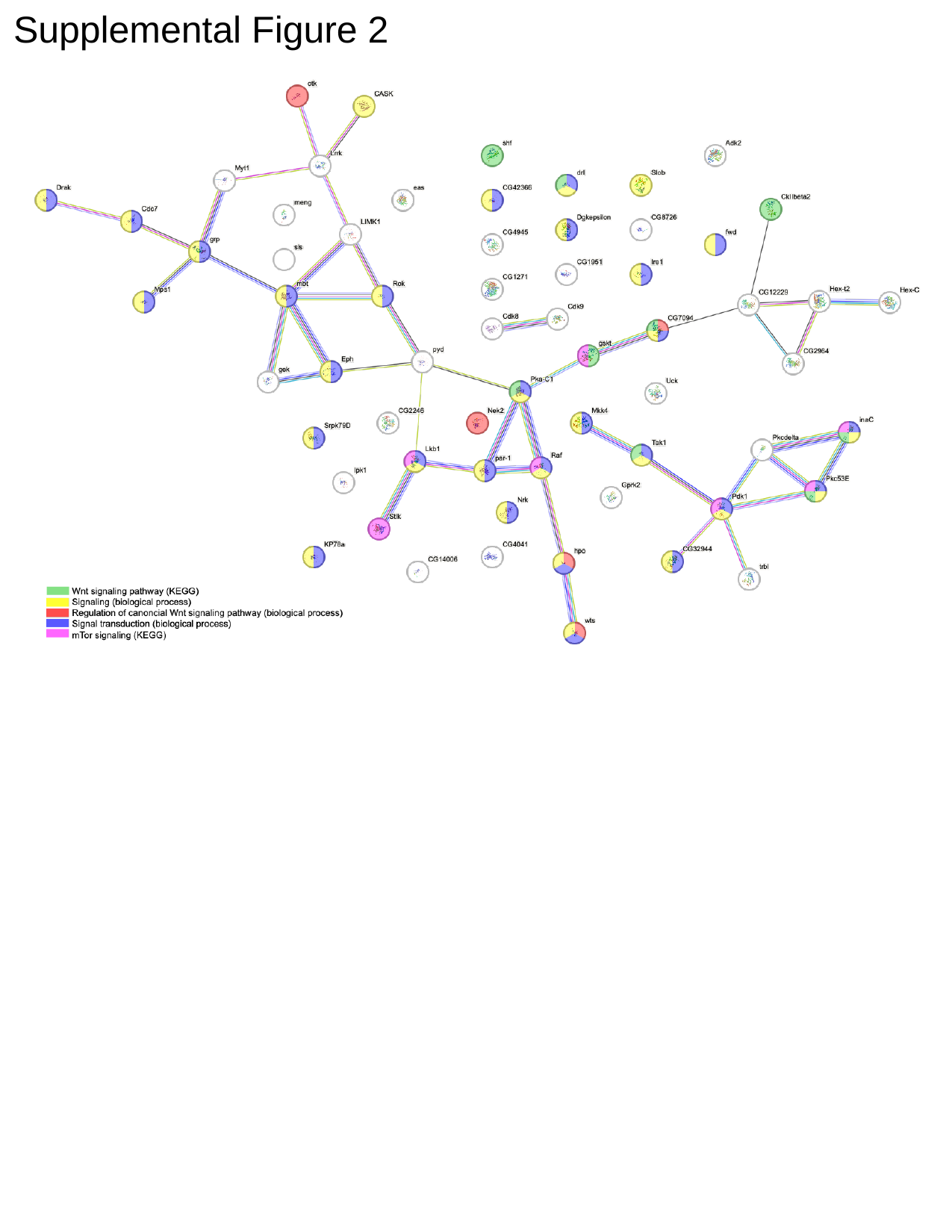

Supplemental Figure 2

### Slide 3
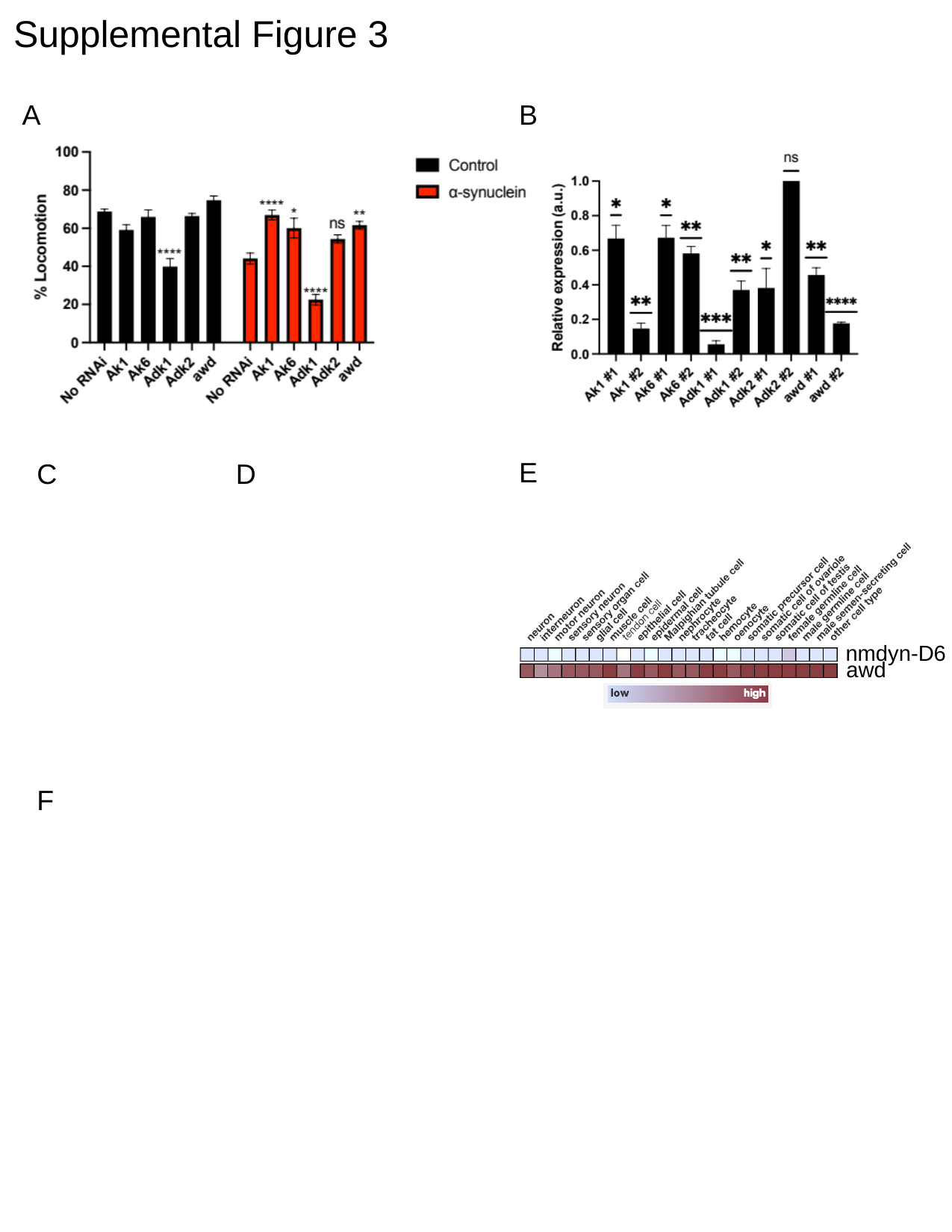

Supplemental Figure 3
A
B
E
C
D
nmdyn-D6
awd
F

### Slide 4
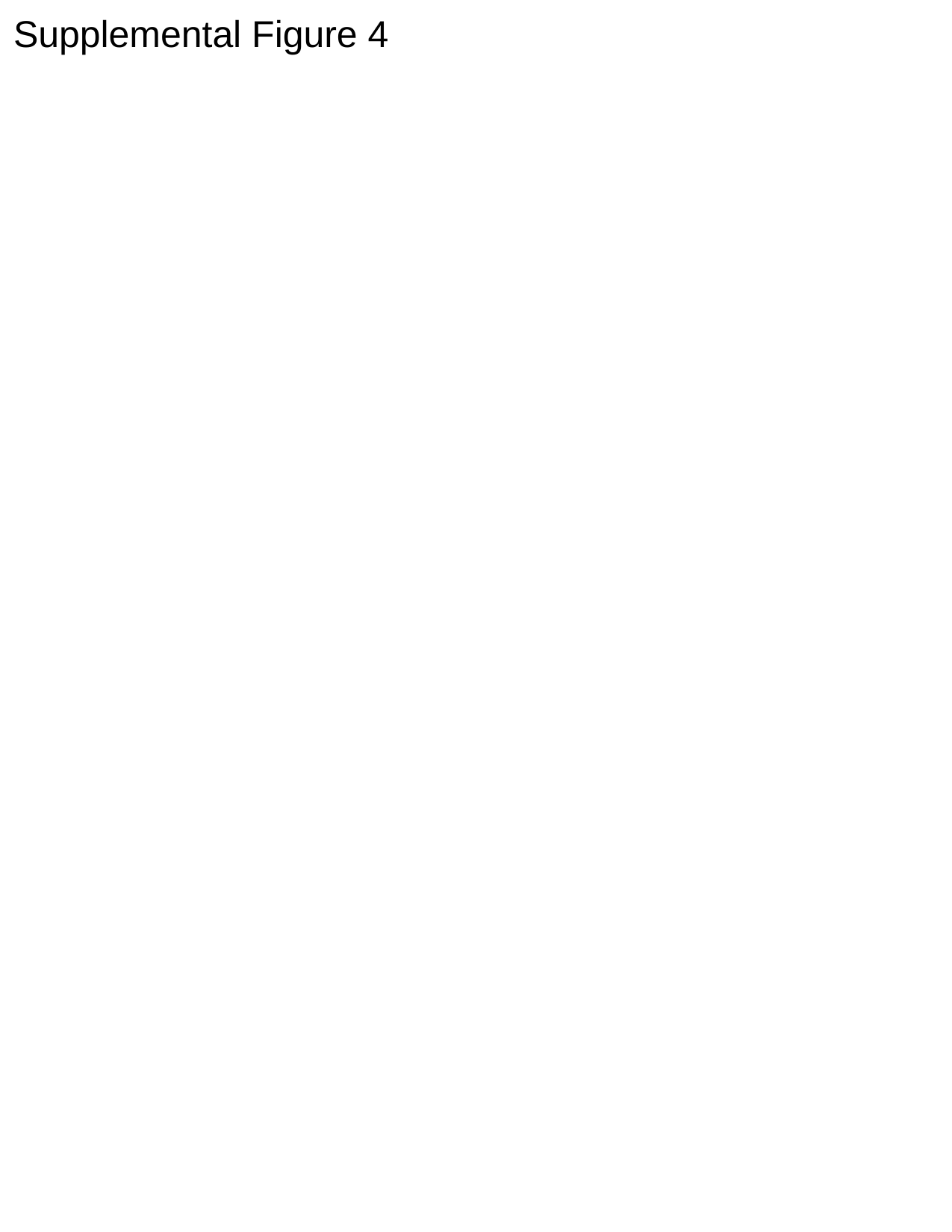

Supplemental Figure 4

### Slide 5
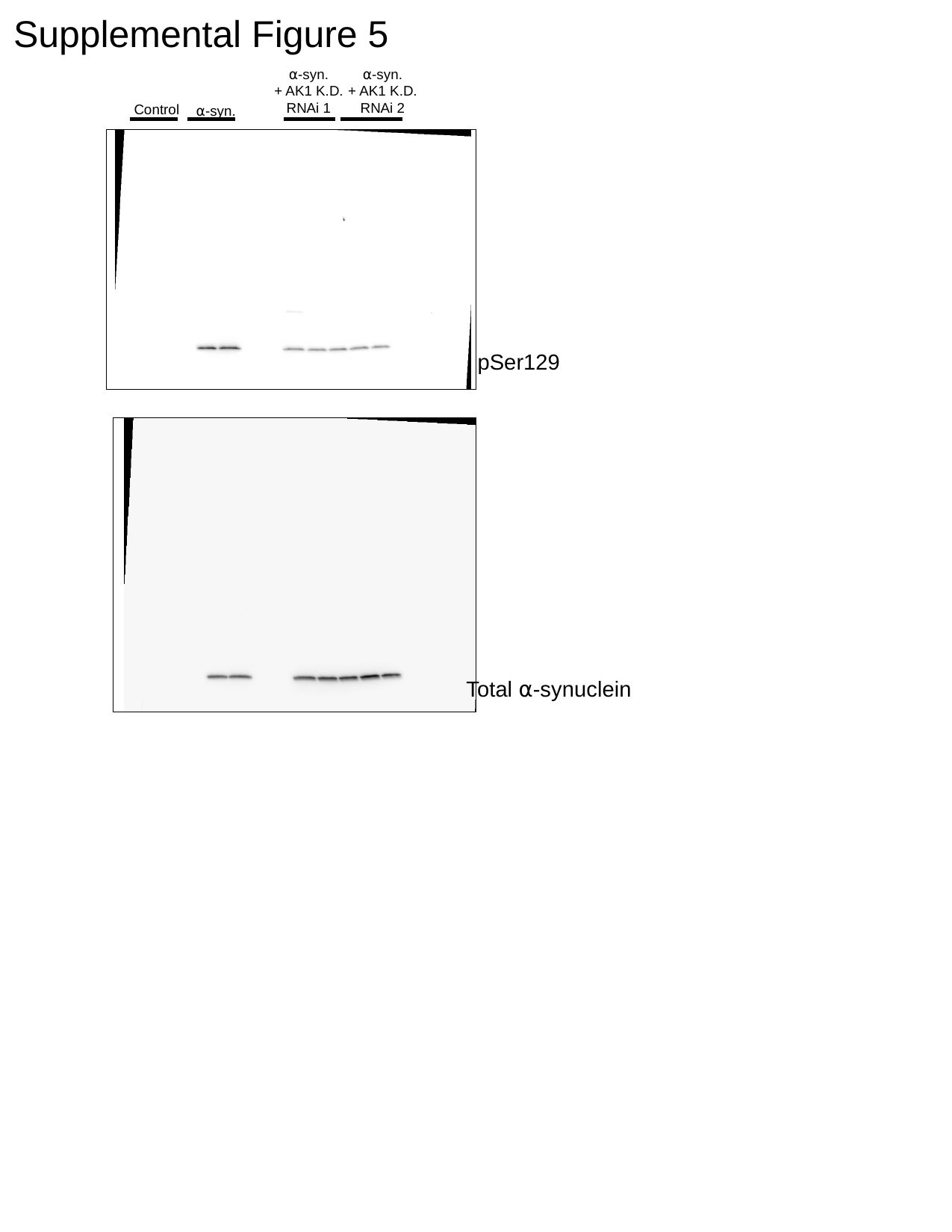

Supplemental Figure 5
⍺-syn.
+ AK1 K.D.
RNAi 2
⍺-syn.
+ AK1 K.D.
RNAi 1
Control
⍺-syn.
pSer129
Total ⍺-synuclein
